# Refining the Application of Microbial Lipids as Tracers of *Staphylococcus aureus* Growth Rates in Cystic Fibrosis Sputum

**DOI:** 10.1101/348037

**Authors:** Cajetan Neubauer, Ajay S. Kasi, Nora Grahl, Alex L. Sessions, Sebastian H. Kopf, Roberta Kato, Deborah A. Hogan, Dianne K. Newman

## Abstract

Chronic lung infections in cystic fibrosis (CF) could be treated more effectively if the effect of antimicrobials on pathogens *in situ* were known. Here, we compared changes in the microbial community composition and pathogen growth rates in longitudinal studies of CF patients undergoing intravenous antibiotic administration during pulmonary exacerbations. Microbial community composition was measured by NanoString DNA analysis and growth rates were obtained by incubating CF sputum with heavy water and tracing incorporation of deuterium into two different *anteiso* fatty acids (*a*-C_15:0_ and *a*-C_17:0_) using gas chromatography–mass spectrometry (GC/MS). Prior to this study, both lipids were thought to be specific for Staphylococcaceae and hence their isotopic enrichment was interpreted as a growth proxy for *S. aureus.* Our experiments revealed, however, that *Prevotella* is also a relevant microbial producer of *a*-C_17:0_ fatty acid in some CF patients, thus deuterium incorporation into these lipids is better interpreted as a more general pathogen growth rate proxy. Even accounting for a small non-microbial background source detected in some patient samples, *a*-C_15:0_ fatty acid still appear to be a relatively robust proxy for CF pathogens, revealing a median generation time of ~1.5 days, similar to prior observations. Contrary to our expectations, pathogen growth rates remained relatively stable throughout exacerbation treatment. We suggest two best practices for application of stable isotope probing in CF sputum: (1) parallel determination of microbial community composition in CF sputum using culture-independent tools, and (2) analysis of samples with a minimum *a*-C_15:0_ concentration of 0.1 weight percent of saturated fatty acids.

**IMPORTANCE:** In chronic lung infections, populations of microbial pathogens change and mature in ways that are often unknown, which makes it challenging to identify appropriate treatment options. A promising tool to better understand the physiology of microorganisms in a patient is stable-isotope probing, which we previously developed to estimate the growth rates of *S. aureus* in cystic fibrosis (CF) sputum. Here, we tracked microbial communities in a cohort of CF patients and found that *anteiso* fatty acids can also originate from other sources in CF sputum. This awareness led us to develop an new workflow for the application of stable isotope probing in this context, improving our ability to estimate pathogen generation times in clinical samples.

## INTRODUCTION

Cystic Fibrosis (CF) is a genetic disease that affects more than 30,000 individuals in North America with a median predicted survival age of 41.6 years (1). This disease is caused by mutations in the CF transmembrane conductance regulator (CFTR) protein that functions as an apical epithelial chloride channel. Reduced or absent chloride channel function leads to dehydrated, viscous secretions that cause dysfunction in the lungs, pancreas and other organs. As CF progresses, bacteria colonize the lung, resulting in chronic polymicrobial infections and inflammation (2). These infections are notoriously hard to treat because species, strains within a species, and genomic variants within a strain differ between individuals and discrete regions of the lung; infections become more resistant to drug treatment as the pulmonary disease progresses (3, 4). Understanding which pathogens are present can inform the choice of appropriate treatment strategies (5) and improve lung function, in tandem with restoration of CFTR activity by CFTR potentiator drugs such as Ivacaftor (6).

*Staphylococcus aureus* is one of the earliest bacteria detected in infants and children with CF (7). The development of chronic polymicrobial infections in the CF lung involves many contributing factors. *S. aureus* is the most prevalent bacterium (70.6%) detected on culture in patients with CF in the US (1). Over the past decade, the prevalence of methicillin-resistant *Staphylococcus aureus* (MRSA) has increased in CF patients in the United States. Compared to CF patients with methicillin-sensitive *S. aureus* (MSSA) infections, CF patients with MRSA have lower lung function, increased likelihood of hospitalization and lack of responsiveness to antibiotic treatment (8). Persistent MRSA infection in CF patients between 8-21 years is associated with more rapid lung function decline (9).

CF patients experience episodes of acute worsening of respiratory symptoms called pulmonary exacerbations. Pulmonary exacerbations are treated with antibiotics and airway clearance therapies (10). For decades, antibiotics have been the cornerstone of CF care and in recent years, there has been a clinical focus on the aggressive management of pulmonary exacerbations because 25% of CF patients do not recover their baseline lung function after intravenous antibiotic treatment for a pulmonary exacerbation (11). A higher risk of failing to recover to baseline lung function was associated with several factors, including persistent infection with *P. aeruginosa* or MRSA (11).

A largely unresolved question is how quickly CF pathogens proliferate in the lung during antibiotic therapy (12–15). It is conceivable that the ability of infections to persist in the lung is connected in intricate ways to slow growth rates. Slow bacterial growth, for example in biofilms, can be beneficial for survival partly because many drugs have their greatest effect on rapidly dividing cells (16). Nevertheless, for a microbial population to persist in the lung, a certain amount of growth is necessary to counteract loss of cells due to attack by the immune system or mucociliary clearance. As chronic infections have highly complex and heterogeneous composition, it is challenging to determine *in situ* growth rates of CF pathogens.

Recently, two advances in microbial ecology have enabled to measure microbial growth rates *in situ*. The first is based on DNA sequencing and exploits the observation that growing cells yield more sequencing reads at genomic regions near the origin of replication (17, 18). This difference between non-growing and rapidly growing cells is relatively small (a maximum 2-fold increase when assuming duplication of origin of replication in growing cells). To derive quantitative information from this signal, metagenomic data must have high sequence coverage, assumptions must be made about chromosome copy number in growing cells, and care must be taken to avoid biases introduced by preferential DNA sequencing from certain members of the population. The second approach uses isotopic labeling and mass spectrometry to quantify the incorporation of isotopes into biomass, which provides a more direct measurement of anabolic activity (14, 19). Stable isotope probing requires incubation of the sample with isotopic label and relies on the identification of diagnostic metabolites. Because the natural abundance of rare isotopes such as deuterium is small and the number of metabolite molecules per cell is large, the sensitive detection of isotope incorporation into metabolites by mass spectrometry can have a greater dynamic range than DNA sequencing. Thus, stable-isotope probing may be better suited for studies that attempt to gain growth information for microbes in a sample where microbes would be expected to be growing slowly and the DNA of the target organism(s) might only make up only a small portion of the total DNA. While DNA sequencing and stable isotope probing provide new opportunities to study *in situ* growth rates, both methods provide information by analyzing bulk extracts and cannot resolve spatial organization and heterogeneity (14), though techniques are emerging that permit microbial metabolic activity to be measured at the single-cell level (20–22).

Utilizing a bulk stable-isotope labeling approach, we previously attempted to quantify *S. aureus* growth rates in CF sputum; we found that its growth rate was slowest in acutely sick patients, despite extensive growth rate heterogeneity at the single-cell level (14). Freshly expectorated sputum was incubated with heavy water (D_2_O) and incorporation of deuterium into diagnostic fatty acids was measured. These fatty acids, *anteiso*-C15:0 (*a*-C_15:0_) and *anteiso*-C17:0 (*a*-C_17:0_), are the dominant fatty acids of *S. aureus* and their biosynthesis rate was used as a proxy for growth of the pathogen. The two main findings of the previous study (14) were: (1) *S. aureus* had a median *in situ* generation time of ~2 days, far slower than previous estimates of CF pathogen growth rates, and (2) growth rates were slowest in patients during early treatment of pulmonary exacerbations (*i.e.* first few days). The slow growth rates we measured support the hypothesis that *S. aureus* populations experience environmental constraints within the lung, which could facilitate tolerance to conventional antibiotics. Based on the observed cross-sectional growth trends (14), we hypothesized that during early hospitalization, treatment with anti-Staphylococcal antibiotics may have suppressed *S. aureus* growth rates but as other members of the microbial community also succumbed to antibiotic treatment, *S. aureus* could occupy these abandoned niches and grow more quickly.

This study was designed to test this hypothesis by performing a longitudinal study. We correlated changes of microbial community composition with *S. aureus* growth rate in a cohort of CF patients that were treated with intravenous antibiotics due to pulmonary exacerbations. Our results did not support our hypothesis, but lead to important refinements in our approach to measuring *in situ* growth rates in clinical samples.

## RESULTS

We collected and analyzed 8 time series of sputum samples during pulmonary exacerbations from seven CF patients. Three or more longitudinal samples were acquired from all patients during their hospitalization. When possible, a sputum sample was collected during a routine CF clinic visit three to four weeks following hospitalization. All enrolled patients were positive for *S. aureus* based on clinical microbiology culturing procedures. Two patients had severe lung disease (Patient 1 and 7), the remaining longitudinal series were collected from patients with mild to moderate lung disease. All patients recovered to their spirometric baseline forced expiratory volume in one second (FEV1%) following treatment. Clinical and demographic information for study participants is summarized in Table 1 (see also Table S1 and S2).

The individual time series revealed dramatic changes in the microbial populations based on counting ribosomal RNA within total RNA extracted from sputum using NanoString technology (Figure 1). Though for this study we specifically recruited CF patients with *S. aureus* infections on the basis of clinical cultures, polymicrobial infections were the norm, as expected (23). In three patients, *S. aureus* was the most abundant member of the microbial community at the beginning of treatment (Patient 1, 2, 3). In some patients *Prevotella melaninogenica* (Patient 4, 5) or *Pseudomonas aeruginosa* (Patient 6, 7) were most abundant. For Patient 7, *P. aeruginosa* accounted for almost all (>96%) of NanoString counts, while *S. aureus* was detected only at the beginning of hospitalization (0.4% abundance). During treatment, the relative abundance of *S. aureus* decreased in most patients as assessed by rRNA levels. Patient 1, however, who had severe lung disease, showed a varied response. Based on total NanoString counts we can infer microbial population size (Figure 1) (24). Several patients showed a large decrease in microbial population size throughout their hospital stay (Patients 2, 3, 5 and 6), whereas in Patient 1, microbial population size did not decrease, and even increased in Patients 4 and 7. These time series illustrate that CF microbial infections change and mature differently in individuals during antimicrobial therapy. Accordingly, we might expect that differential microbial growth activities during drug treatment in any given patient would correlate with the rise or fall of particular members of the microbial community.

**Figure 1.**
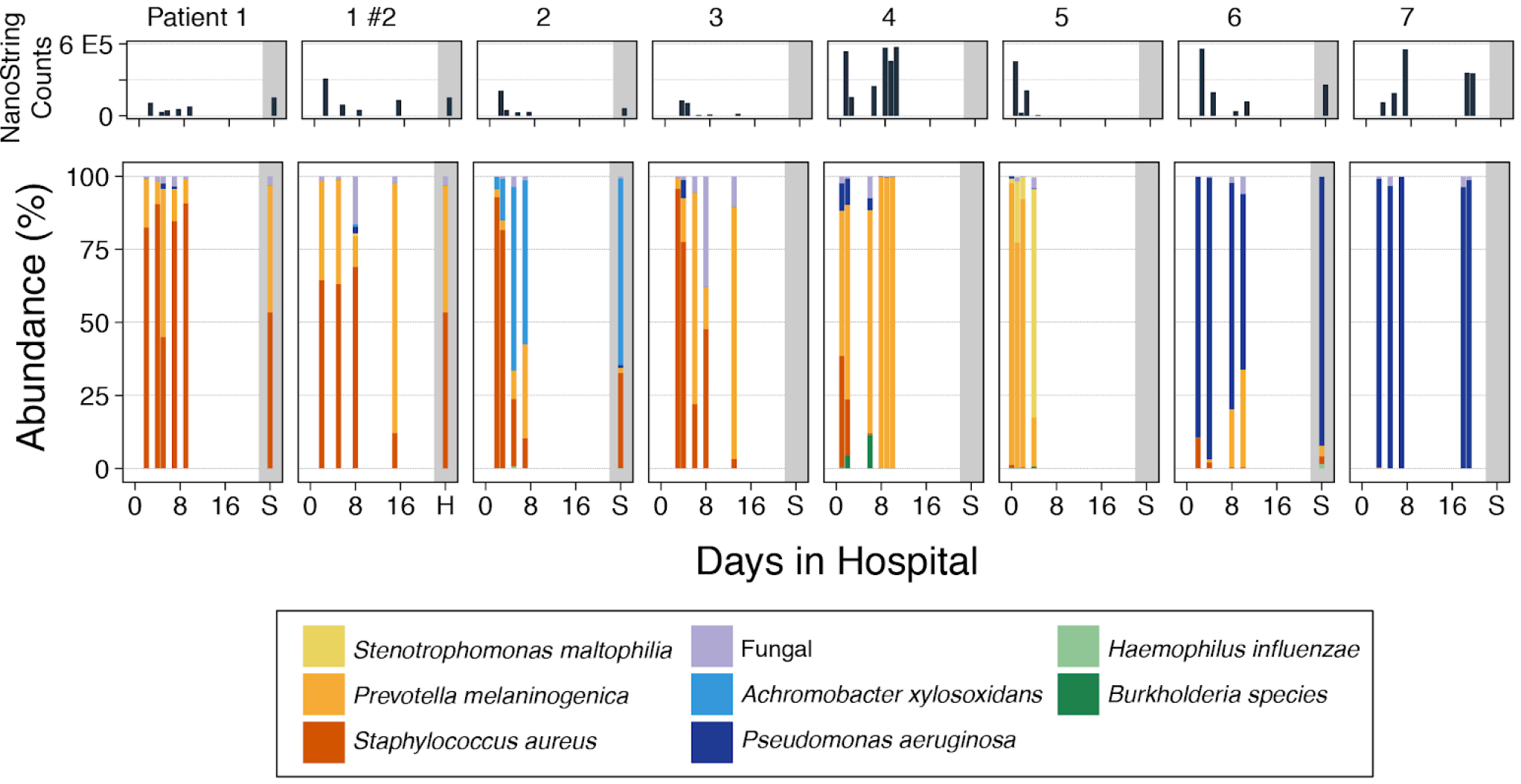
Microbial community composition of expectorated CF sputum during pulmonary exacerbations. Expectorated CF sputum from eight time courses was used to isolate RNA for counting of rRNA molecules by NanoString, using CF-specific probes (see Methods). Patients were hospitalized and treated with intravenous antibiotics. For each longitudinal series the top panel shows the sum of microbial rRNA counts, an indicator of microbial abundance. The bottom panel shows the relative abundance of common CF-associated microorganisms. Producers of *anteiso* fatty acids are colored yellow to red. S: Control sample (‘stable’) taken at a routine visit to the clinic after treatment.

To determine whether this was the case, we focused on measuring the growth activity of *S. aureus* by tracking *anteiso* fatty acids previously thought to be diagnostic for this bacterium in the context of CF microbial communities, comparing its specific growth rates to changes in the microbial community composition (Figure 2). Although *S.aureus* was detected in initial sputum cultures of all patients included in the longitudinal study, our analysis of microbial populations showed that Patient 7 contained almost exclusively *P. aeruginosa*. This bacterium does not produce *anteiso* fatty acids so we did not expect to detect *a*-C_15:0_ and *a*-C_17:0_ analytes in samples from Patient 7. Nevertheless, *a*-C_17:0_ occurred at abundances >0.15% (wt% of saturated fatty acids), which is similar to the levels in sputum from patients that had dominant *S. aureus* infections (e.g., Patient 1). For comparison, we analyzed sputum samples from three CF patients whose clinical data did not indicate presence of *S. aureus*. Two of these controls also contained more than 0.15% *a*-C_17:0_ in two samples. Interestingly, *a*-C_15:0_ was detected at lower abundances (<0.05%) in Patient 7 and the three controls. This suggested that sources other than *S. aureus* can contribute *anteiso* fatty acids, especially *a*-C_17 0_, in expectorated sputum.

**Figure 2.**
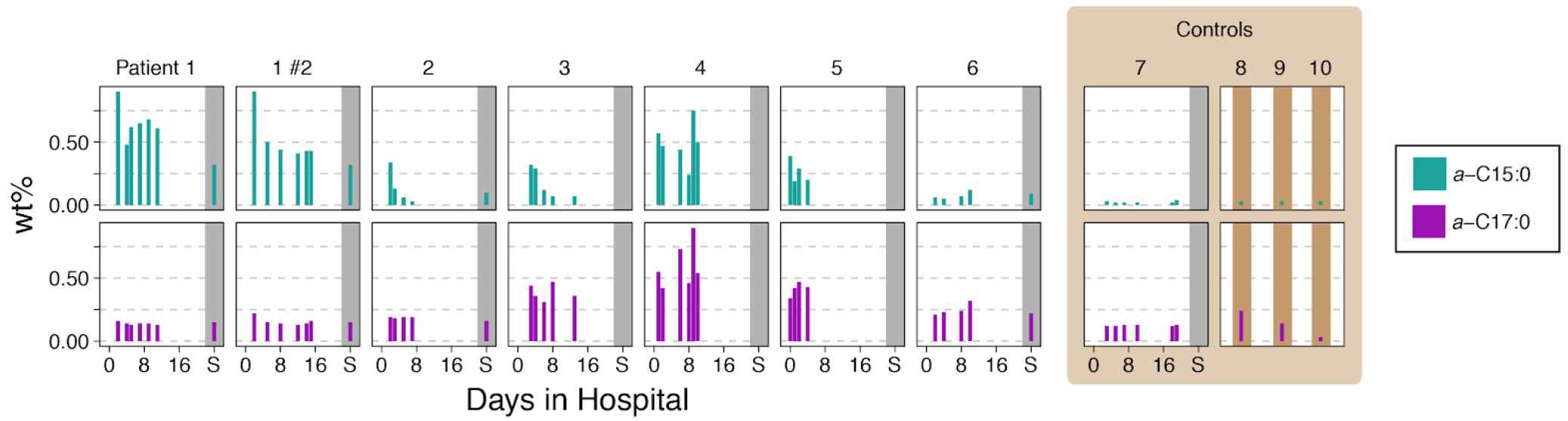
Abundance of *anteiso* fatty acids in expectorated CF-sputum. The wt% abundance of *a*-C_15:0_ (cyan) and *a*-C_17:0_ (magenta) in extracts of saturated fatty acids are shown. Patient 7, whose sputum did not contain common microbial producers of *anteiso* fatty acids, is grouped together with control samples that were collected from three individuals whose clinical data did not indicate presence of *S. aureus.*

Whether non Staphylococcal sources of *anteiso* fatty acids were sufficiently abundant to affect the interpretation of our D_2_O-labeling experiments became the next question. Our prior work had taught us that *Stenotrophomonas* species can also produce *anteiso* fatty acids and can occur in CF sputum (14). In some samples from Patient 5, *Stenotrophomonas maltophilia* constituted the majority of microorganisms. The fatty acid analysis of these sputum samples also revealed an increased amount of *iso*-C_15:0_, which had been observed for the pure strain (14). In all other patients, *S. maltophilia* occurred at low abundances (<1%), too low to account for the high levels of *a*-C_170_ we measured. For example, in Patient 4 neither *S. aureus* nor *S. maltophilia* were detected, yet *a*-C_15:0_ and *a*-C_17:0_ were both present at up to 0.8% of saturated fatty acids. A likely source of *anteiso* fatty acids in sputum from Patient 4, and samples from other patients in our cohort, is *Prevotella melaninogenica*. This species can produce *anteiso* fatty acids but it was not previously recognized as a relevant source of *anteiso* fatty acids in CF sputum (14, 25). *P. melaninogenica* is an anaerobe and part of the oral flora. It was isolated from CF sputum and in bronchoalveolar lavage fluid from infants with CF (26, 27), yet the clinical significance of *Prevotella* in CF lung disease is still debated (28).

Sputum from Patient 7 had elevated *a*-C_170_ but did not contain *Prevotella* (Figure 2). This raised the new concern that perhaps additional input of *anteiso* fatty acids might emanate from the human host itself. *Anteiso* fatty acids are rare in human tissues but can accumulate in skin and fat cells (29). Might human cells, such as neutrophils and epithelial cells, contribute to the *anteiso* fatty acids pool in expectorated CF sputum? To answer this question, we analyzed the fatty acid composition of two human cell lines (Table 2). CF bronchial epithelial cells (CFBE) and polymorphonuclear neutrophils (PMN) indeed contained *anteiso* fatty acids on the order of 0.1 wt% of total fatty acids. It is possible that some of these *anteiso* lipids in cell cultures originate from the growth substrate. Fetal bovine serum (FBS), the only component of the growth medium that contains lipids, contained a similar percentage of *anteiso* fatty acids and hence diet could also be a contributing factor. For humans, dietary intake of *anteiso* fatty acids in the United States is largely via milk and meat products (30). We concluded that an additional source of *anteiso* fatty acids in expectorated sputum likely stems from human cells or dietary intake. The impact of this contribution on the interpretation of D_2_O-labeling experiments will depend on the relative contribution of human cells to the pool of *anteiso* fatty acids in a given sample.

**Table 1.**
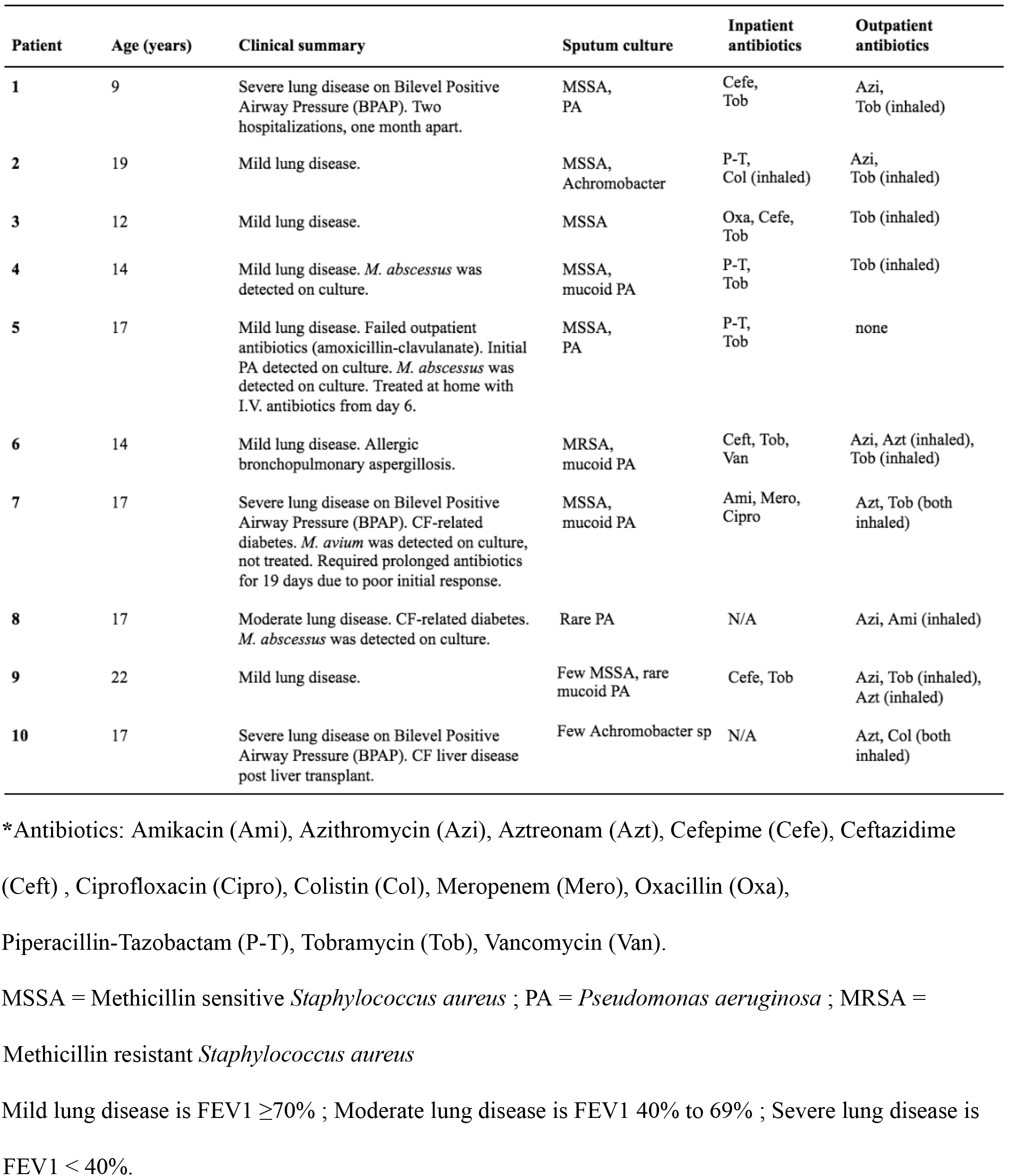
Clinical summary

| Patient | Age (years) | Clinical summary | Sputum culture | Inpatient antibiotics | Outpatient antibiotics |
| --- | --- | --- | --- | --- | --- |
| 1 | 9 | Severe lung disease on Bilevel Positive Airway Pressure (BPAP). Two hospitalizations, one month apart. | MSSA, PA | Cefe, Tob | Azi, Tob (inhaled) |
| 2 | 19 | Mild lung disease. | MSSA, Achromobacter | P-T, Col (inhaled) | Azi, Tob (inhaled) |
| 3 | 12 | Mild lung disease. | MSSA | Oxa, Cefe, Tob | Tob (inhaled) |
| 4 | 14 | Mild lung disease. *M. abscessus* was detected on culture. | MSSA, mucoid PA | P-T, Tob | Tob (inhaled) |
| 5 | 17 | Mild lung disease. Failed outpatient antibiotics (amoxicillin-clavulanate). Initial PA detected on culture. *M. abscessus* was detected on culture. Treated at home with I.V. antibiotics from day 6. | MSSA, PA | P-T, Tob | none |
| 6 | 14 | Mild lung disease. Allergic bronchopulmonary aspergillosis. | MRSA, mucoid PA | Ceft, Tob, Van | Azi, Azt (inhaled), Tob (inhaled) |
| 7 | 17 | Severe lung disease on Bilevel Positive Airway Pressure (BPAP). CF-related diabetes. *M. avium* was detected on culture, not treated. Required prolonged antibiotics for 19 days due to poor initial response. | MSSA, mucoid PA | Ami, Mero, Cipro | Azt, Tob (both inhaled) |
| 8 | 17 | Moderate lung disease. CF-related diabetes. *M. abscessus* was detected on culture. | Rare PA | N/A | Azi, Ami (inhaled) |
| 9 | 22 | Mild lung disease. | Few MSSA, rare mucoid PA | Cefe, Tob | Azi, Tob (inhaled), Azt (inhaled) |
| 10 | 17 | Severe lung disease on Bilevel Positive Airway Pressure (BPAP). CF liver disease post liver transplant. | Few Achromobacter sp | N/A | Azt, Col (both inhaled) |

\*Antibiotics: Amikacin (Ami), Azithromycin (Azi), Aztreonam (Azt), Cefepime (Cefe), Ceftazidime
(Ceft) , Ciprofloxacin (Cipro), Colistin (Col), Meropenem (Mero), Oxacillin (Oxa),
Piperacillin-Tazobactam (P-T), Tobramycin (Tob), Vancomycin (Van).
MSSA = Methicillin sensitive *Staphylococcus aureus* ; PA = *Pseudomonas aeruginosa* ; MRSA =
Methicillin resistant *Staphylococcus aureus*
Mild lung disease is FEV1 $\geq 70\%$ ; Moderate lung disease is FEV1 40% to 69% ; Severe lung disease is
FEV1 $< 40\%$ .

**Table 2.** Concentrations of *anteiso* fatty acids in various sources (wt% of fatty acids)

| Source | *a*-C<sub>15:0</sub> | *a*-C<sub>17:0</sub> | Reference |
| --- | --- | --- | --- |
| *S. maltophilia* | 20 | <1 | Kopf *et al.* |
| *S. aureus* | 45 | 20 | Kopf *et al.* |
| *P. melaninogenica* | 43 | 4 | Wu *et al.* |
| *S. epidermidis* | 28 | 20 | This study |
| CF bronchial epithelial cells (CFBE) | 0.05 | 0.3 | This study |
| Human promyelocytic leukemia cells (HL-60) | 0.04 | 0.1 | This study |
| Polymorphonuclear neutrophils (PMN) | 0.07 | 0.1 | This study |
| Fetal bovine serum (FBS) | 0.04 | 0.3 | This study |
| Diet | 0.23 (beef)<br>0.56 (milk) | 0.54 (beef)<br>0.61 (milk) | Ran-Ressler *et al.* |

Given these new findings, what can we say with confidence about the growth rate of any particular pathogen in sputum? The deuterium labeling of *anteiso* fatty acids, interpreted at face value, yielded a median generation time of *S. aureus* of 2.0 days when *a*-C_150_ is used as proxy for microbial growth. The longer anteiso fatty acid, *a*-C_17:0_, yielded systematically slower generation time with a median of 14.6 days. The individual time series in our data set reveal a large variation between patients when deuterium incorporation is interpreted without taking microbial abundance and community composition into account (Figure 3). For example, the second time series for Patient 1 (1 #2), who had two hospitalizations one month apart, showed a slower growth rate during hospitalization followed by a rebound increase in growth rate. This patient had severe lung disease (baseline FEV1 34%) requiring non-invasive ventilation during sleep and treatment for chronic airway *P. aeruginosa* colonization with cycled nebulized tobramycin. However, a similar growth rate trend was not observed in the same patient during the first hospitalization (Patient 1 #1) and in other patients. It is possible that these discrepancies could be due to stochastic differences in sputum composition with respect to the proportion of human:microbial cells at these different time points. Whether or not this was the case, the majority of our longitudinal time series did not show an obvious connection between the duration of intravenous antibiotic treatment and deuterium labeling of *anteiso* fatty acids.

**Figure 3.**
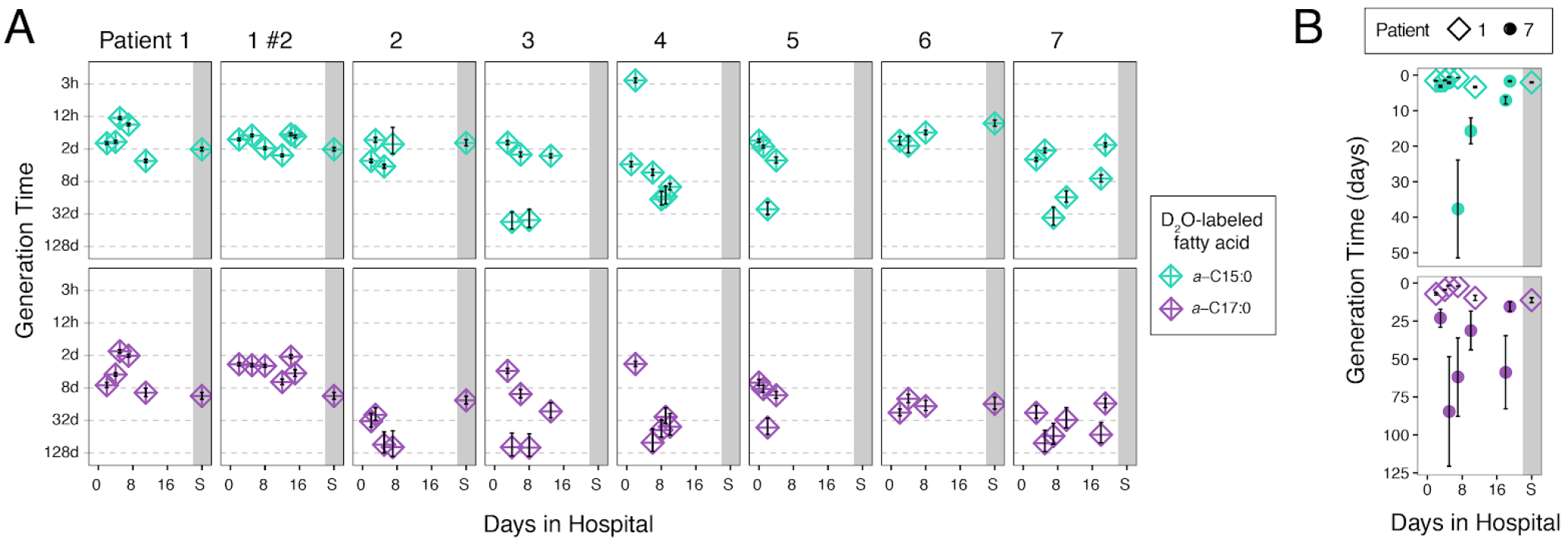
Generation times based on deuterium incorporation into *anteiso* fatty acids. **A.** Generation time estimates obtained for the eight time series. Error bars represent ±1SD. **B.** Data for Patient 1 (open diamonds), which had abundant microbial producers of *anteiso* fatty acids, and Patient 7 (closed circles), which had no detectable microbial producers of anteiso fatty acids, are plotted on a linear scale for better comparison.

Most samples in our longitudinal dataset contain *a*-C_15:0_ at concentrations that are significantly higher (10 to 50-fold) than those of Patient 7, in which *S. aureus* is absent (Figure 2). Accordingly, *a*-C_15:0_ in these samples is likely providing information about the growth of other CF-associated microbes that produce *a*-C_15:0_, namely *P. melaninogenica* and *S. maltophilia.* Whether isotope incorporation yields reliable growth rates for any pathogen also depends on the labeling rates of *anteiso* fatty acids from non-microbial origins. Non-microbial sources mainly appear to contribute *a*-C_17:0_, which in CF sputum generally shows slower label incorporation than *a*-C_15:0_ (14). Accordingly, contributions of non-microbial *anteiso* fatty acids tend to decrease growth rate estimates.

To better constrain how non-microbial *anteiso* fatty acids might affect our ability to infer microbial growth rates, we analyzed their potential impact using modeling. To a first approximation, we assume the pool of *anteiso* fatty acids is a mixture of two end members at the beginning of the labeling experiment: a portion of biosynthetically active microbial lipids (*x*) and a portion of inactive dietary lipids (*1-x*). Assuming that bacterial fatty acids are produced exponentially at a constant generation time (*GT*) and dietary fatty acids remain unlabeled and constant during incubation with D_2_O, we can calculate the apparent generation time (*y*) using:

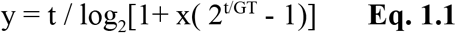

The effect of non-microbial sources of *anteiso* fatty acids on bacterial growth rate estimates from this basic model is shown in Figure 4A. For example, if the microbes have an actual generation time of 2 days and 90% of the *anteiso* fatty acids originate are microbial origin and the remaining 10 % dietary, the apparent generation time after 1 hour of incubation would be 2.2 days, *i.e.* only slightly slower than the actual doubling every 2 days.

**Figure 4.**
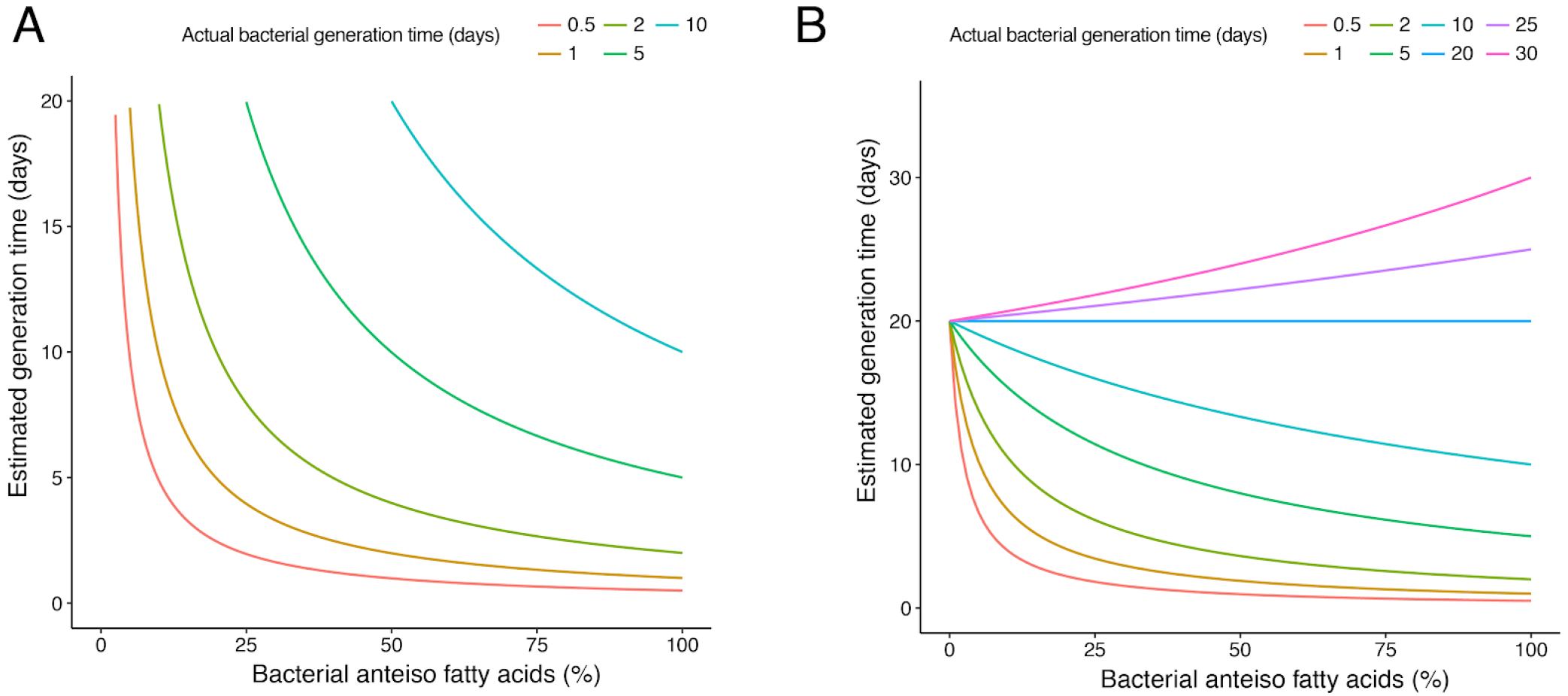
Effect of non-microbial sources of *anteiso* fatty acids on bacterial growth rate estimates. **A.** The sensitivity analysis assumes exponential production of isotopically labeled *anteiso* lipids for different doubling times (colored lines) and zero isotope incorporation into *anteiso* fatty acids from other sources, e.g. diet. Equation Eq. 1.1 was used to calculate generation times. **B.** Same as in first panel, but assuming that the non-microbial fraction of the *anteiso* fatty acid pool has isotope incorporation with a doubling time of 20 days.

We can extend this basic model to include scenarios in which non-bacterial fatty acids can become isotopically labeled, e.g. due to *de novo* biosynthesis in human cells. If we assume a constant generation time *GT*_*b*_ for the bacterial fraction (*x*) and *GT*_*n*_ for the non-bacterial fraction (*1-x*), the apparent generation time *y* is given by:

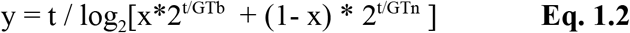

Figure 4B shows the effect of non-bacterial growth with a generation time of 20 days (*GT*_*n*_), which is on the order of generation times measured for *a*-C_17:0_ in samples from Patient 7 (presumed to be non-bacterial in origin). In this extended model, the non-bacterial fatty acids have a relatively slow but non-zero isotope incorporation, as a result the deviations between actual and apparent microbial growth rates tend to be smaller than the simpler model.

This sensitivity analysis suggests that *a*-C_15:0_ can be interpreted as a robust proxy for bacterial growth in CF infections provided most of the analyte originates from bacteria. In sputum from Patient 7, *a*-C_15:0_ did not derive from any known microbial sources and was detected at a mean concentration of 0.025 wt%. Using 4-fold this value (0.1 wt%) as a threshold, we filtered our longitudinal data to identify samples for which microbial growth rates in sputum could be obtained reliably from deuterium labeling of *a*-C_15:0_. The median growth rates estimated for these 23 samples is 1.9 days, which represents a mixture of growth rates for all three *anteiso* fatty acid producing microbial strains (Table 3). As expected, the amount of *anteiso* fatty acids detected in CF sputum has a moderately positive correlation with the amount of microbial producers (Figure S1; Pearson correlation coefficient +0.52). This analysis suggests that *P. melaninogenica* and *S. aureus* are the predominant microbial producers of *anteiso* fatty acids in most expectorated CF sputum samples in our cohort. When we compared the abundance of microbial strains with the generation times in the 23 samples with high *a*-C_15:0_ content (Table 3), we found that samples that were dominated by *P. melaninogenica* (median 2.5 days; 11 samples) and *S. maltophilia* (3.2 days; 1 sample) had slower generation times than those which contains >50% of *S. aureus* (1.5 days; 11 samples). Based on this refined analysis of *anteiso* fatty acids, we did not observe a consistent decrease of bacterial growth rates during pulmonary exacerbation treatment with intravenous antibiotics in individual patients. The only *S. aureus-rich* sample that had a much slower growth was from Patient 3 (generation time 45 days; day 4). By the following time point (day 6), the bacterial community size had dropped by 94 % and the relative abundance of *S. aureus* had decreased from 78% to 22%, while growth recovered to a generation time of 2.5 days. This indicates that low levels of label incorporation into *a*-C_15:0_ can coincide with recent population decline. Killed *S. aureus*, including intact phospholipids, may transiently remain in sputum and cause slow apparent growth rates independent of the growth rate of surviving bacteria. In summary, *S. aureus* generation time in our cohort was typically on the order of 1-2 days and did not consistently decrease during antibiotic treatment of pulmonary exacerbations.

**Table 3.** Summary of samples for which growth rates can be reliably inferred from *a*-C_15:0_

| Patient | Day | α-C15:0<br>(wt%) | GT [days] | *S. aureus*<br>(%) | *Stenotrophomonas*<br>(%) | *Prevotella* (%) |
| --- | --- | --- | --- | --- | --- | --- |
| 1 | 2 | 0.90 | 1.6 | 82.5 | 0 | 16.8 |
| 1 | 4 | 0.48 | 1.5 | 90.5 | 0 | 7.8 |
| 1 | 5 | 0.62 | 0.5 | 45.0 | 0 | 50.8 |
| 1 | 7 | 0.65 | 0.7 | 84.7 | 0 | 11.1 |
| 1 | Stable | 0.32 | 2.0 | 53.4 | 0 | 43.5 |
| 1 #2 | 2 | 0.90 | 1.3 | 64.5 | 0 | 34.0 |
| 1 #2 | 5 | 0.50 | 1.1 | 63.1 | 0 | 35.7 |
| 1 #2 | 8 | 0.44 | 1.9 | 69.1 | 0.9 | 10.5 |
| 1 #2 | 15 | 0.43 | 1.2 | 12.2 | 0 | 85.6 |
| 2 | 2 | 0.34 | 3.3 | 92.8 | 0 | 2.8 |
| 2 | 3 | 0.13 | 1.4 | 81.6 | 0 | 3.3 |
| 3 | 3 | 0.32 | 1.5 | 95.8 | 0 | 3.6 |
| 3 | 4 | 0.29 | 45 | 77.5 | 0 | 15.0 |
| 3 | 6 | 0.12 | 2.5 | 22.1 | 0 | 72.4 |
| 4 | 2 | 0.47 | 0.1 | 19.2 | 0 | 66.6 |
| 4 | 6 | 0.44 | 5.5 | 0.4 | 0 | 76.4 |
| 4 | 8 | 0.24 | 17.1 | 0.1 | 0 | 99.6 |
| 4 | 9 | 0.75 | 15.3 | 0 | 0 | 99.5 |
| 4 | 10 | 0.50 | 10.1 | 0.1 | 0 | 99.7 |
| 5 | 0 | 0.39 | 1.4 | 0.9 | 1.5 | 96.6 |
| 5 | 1 | 0.19 | 1.8 | 0 | 21 | 77.2 |
| 5 | 2 | 0.29 | 26.2 | 0.4 | 7.6 | 91.9 |
| 5 | 4 | 0.20 | 3.2 | 0 | 78.2 | 16.7 |

## DISCUSSION

Our comparison of CF microbiome composition with the abundance of fatty acids indicated that *anteiso* fatty acids in expectorated sputum can originate from previously unrecognized sources, such as *Prevotella* and human cells. Compared to *a*-C_17:0_, lower amounts of *a*-C_15:0_ seem to be introduced by non-microbial sources, making *a*-C_15:0_ a more reliable biomarker for microbial activity. We now view incorporation of deuterium into *a*-C_15:0_ as a likely biomarker for the anabolic activity of *P. melaninogenica* and *S. maltophilia* in addition to *S. aureus.* Fatty acids from human cells or from diet are expected to yield little to no isotope incorporation. From a practical standpoint, best practices for application of deuterium stable isotope probing to CF sputum samples requires knowledge of the microbial community composition and amounts of *a*-C_15:0_ analyte in excess of 0.1 wt% (to be confident that measurements are well above human background levels).

Much of the variability we observe in deuterium labeling data was eliminated when focusing our analysis on samples that had high levels of *a*-C15:0. These samples generally yielded generation times between 0.5 and 2.5 days, with a median of 1.9 days (Table 3). Samples dominated by *S. aureus* (>50%) had a median generation time of 1.5 days. Overall, these estimates for generation times are in the range measured for *a*-C_15:0_ in our previous dataset (14). Because isotopic labeling only provides an average growth estimation for the microbial population, we cannot exclude the possibility that a subpopulation of cells grows significantly slower or faster than with the median generation times we observe. To resolve heterogeneity at this level, single-cell growth rate visualization are necessary (21, 22).

Nevertheless, the comprehensive dataset collected for this longitudinal study allows us to refine our interpretation of whether there are any reproducible patterns in how pathogen growth rates change over the course of antibiotic treatment during a pulmonary exacerbation. After filtering for samples that allow confident estimation of pathogen growth rates (Table 4), we find that growth rate estimates generally remained stable during treatment. We note that samples with low levels of *a*-C_15:0_-producing microorganisms do not yield reliable pathogen growth estimates. However, samples containing high levels of *a*-C_15:0_ and low deuterium incorporation values likely indicate that an antibiotic is inhibiting pathogen growth (Patient 3; day 4). In sum, stable isotope probing in combination with community analysis did not support our previous hypothesis that the *S. aureus* population undergoes niche expansion during antibiotic treatment during an exacerbation. To the contrary, our NanoString data showed that all patients had lower relative abundance of *S. aureus* at the end of treatment compared to the beginning.

In conclusion, stable isotope probing is likely to be most useful in tracking pathogen growth dynamics in response to antibiotic treatment early during pulmonary exacerbations, when pathogenic load is high. With future development, rapid analysis of incubations of CF sputum with different drugs may even help identify the most promising treatment options. Currently, we are developing a method to quantify deuterium label in intact lipids with LC/MS (31). This approach promises to increase the utility of stable isotope probing to assess growth of diverse pathogens.

## METHODS

### Study population

Children and young adults with CF hospitalized for a pulmonary exacerbation were included in this study. Inclusion criteria were a positive diagnosis of cystic fibrosis, presence of *S. aureus* in CF sputum culture, hospitalization for a pulmonary exacerbation, and ability to expectorate sputum. In addition, sputum samples were collected from three CF patients without the presence of *S. aureus* on sputum culture to analyze their fatty acid composition. The study was approved by the Institutional Review Board at Children’s Hospital Los Angeles (IRB# CCI-13-00211). All patients were recruited from Children’s Hospital Los Angeles and informed consent or assent was obtained from all study participants or from a parent/ legal guardian.

The medical records of enrolled subjects were reviewed for age, gender, body mass index, CFTR mutations, duration of hospitalization, results of CF sputum culture, mycobacterial culture, antibiotics used and results of pulmonary function tests. This information is presented in Table 1, S1 and S2.

### Sputum sampling and D_2_O-labeling

Fresh expectorated sputum from children and young adults admitted with a CF pulmonary exacerbation was collected on admission, throughout hospitalization and after return to baseline health. All hospitalized patients received intravenous antibiotics and airway clearance therapies as directed by their treating physician. Expectorated sputum was sampled and treated as in Kopf *et al.* (14). Within 10 minutes of expectoration, sputum samples were segregated into two parts. One part was flash frozen and stored at −80 °C for RNA analysis. The second part (weight > 0.6 g) was incubated at 37 °C for 1 hour with an equivalent weight of prewarmed PBS (phosphate-buffered saline) solution containing 5 % (w/w) D_2_O. The D_2_O content in isotope labelled sputum samples was determined on a DLT-100 liquid water isotope analyzer (Los Gatos Research) (14). To measure the D_2_O content, 150 μL supernatant liquid from the incubated sputum sample was filter-sterilized using a centrifuge tube filter (Costar Spin-X, Corning) and diluted with water of known natural isotopic composition. The remaining sample was flash frozen and stored at −80 °C for fatty acid analysis.

### RNA isolation and analysis

For sputum community profiles, RNA was prepared and analyzed as described by Grahl *et al.* (24). Measurements were made using a NanoString nCounter (NanoString Technologies) (32). We used the custom-designed Nanostring CodeSet described in (24), which enables the simultaneous detection of ribosomal RNA for 38 clinically relevant CF-related bacterial and fungal species or genera. Raw counts were corrected by subtracting 5 times the maximum count for internal negative controls (24).

### Fatty acid extraction, gas chromatography and isotope-ratio measurement

The two target analytes, 14-methyl-hexadecanoic acid (*a*-C_15:0_) and 16-methyl-octadecanoic acid (*a*-C_17:0_), were prepared from sputum samples by transesterification of phospholipids in the presence of a base catalyst (0.5M NaOH in anhydrous methanol), extraction into hexane and clean-up using solid phase extraction (14). Two internal standards (10 μg) were added before extraction (PC(21:0/21:0)) or solid-phase extraction (C_22:0_). The extraction yielded about 10 μg saturated fatty acid methyl esters per 1 mg dry sputum. The fraction containing saturated fatty acid methyl esters (FAMEs) was analyzed by gas chromatography/mass spectrometry (GC/MS) on a Trace DSQ (Thermo Fisher Scientific) with a ZB-5ms column (30 m × 0.25 mm, film thickness 0.25 μm) (14). Peaks were identified by comparison of mass spectra and retention times to authentic standards and library data.

Analytical controls to assess contributions of *a*-C_15:0_ and *a*-C_17:0_ from sources other than *S. aureus* were performed by GC/MS. About 20 mg of lyophilized biomass was transesterified by acid hydrolysis at 100 °C (10 min) in the presence of 20:1 v/v anhydrous methanol/acetyl chloride and hexane (33). Materials tested were *S. epidermidis*, CF bronchial epithelial (CFBE) cells (34), fetal bovine serum (FBS), and polymorphonuclear neutrophils (PMN; differentiated from HL-60 cells).

Deuterium enrichments were quantified in replicates on a DELTA^+^XP (Thermo Fisher Scientific) by GC pyrolysis isotope-ratio mass spectrometry (GC/P/IRMS) as in Kopf *et al.* (14) with the following optimization. The GC program was adjusted for shorter run time and better peak separation of target analytes, *a*-C_15:0_ and *a*-C_17:0_. The GC oven was held at 80 °C for 0.5 min, temperature then was increased at 30 °C/min to 170 °C, held at 170 °C for 45 min, increased again at 30 °C/min to 190 °C, followed by a final increase of 30 °C/min to 320 °C (held for 10 min). Chromatographic peaks were identified in comparison to GC/MS runs by retention order and signal intensity. Reported δD values are corrected for the hydrogen atoms in FAME originating from methanol. Values are stated in the conventional notation versus the VSMOW (Vienna Standard Mean Ocean Water) standard (35). Experimental and analytical errors were estimated as in Kopf *et al.* and we used a Monte Carlo simulation to propagate these uncertainties when calculating growth rate estimates.

## ACKNOWLEDGEMENTS

We thank Elise Cowley, Reto Wijker, Fenfang Wu for helping with our study. We thank Dominique H. Limoli and George O’Toole for providing CFBE cells and Ram Balasubramanian for providing PMN and HL-60 cells. We thank Dr. Jennifer Dien Bard and Dr. Thomas G. Keens for their guidance with our study, the CHLA CF center team and patients of the CHLA CF clinic for participating in this study. We thank the journal editor and reviewers for their comments. This work was funded by grants from the National Institutes of Health (R01HL117328).

